# Functional Yeast Promoter Sequence Design Using Autoregressive Generative Models

**DOI:** 10.1101/2024.10.22.619701

**Authors:** Ibrahim Alsaggaf, Alex A. Freitas, João Pedro de Magalhães, Cen Wan

## Abstract

Functional promoter sequence design plays a crucial role in accurately controlling gene expression processes that are one of the most fundamental mechanisms in biological systems. Thanks to the recent community effort, we are now able to elucidate the associations between yeast promoter sequences and their corresponding expression levels using advanced deep learning methods. This milestone boosts the further development of many downstream biological sequence research tasks including synthetic DNA sequence design. In this work, we propose a novel synthetic promoter sequence generation method, namely Gen-DNA-TCN, which exploits a pre-trained sequence-to-expression predictive model to facilitate its autoregressive generative model training. A large-scale evaluation confirms that Gen-DNA-TCN successfully generates a large number of unique, diverse and functional synthetic yeast promoter sequences that also encode similar transcription factor binding site distributions compared with real yeast promoter sequences.

## 1 Introduction

*Cis*-regulatory mechanisms play a fundamental role in biological systems by allowing transcription factors to interpret regulatory DNA sequences and controlling gene expression levels. A long-standing challenge in biology is to accurately control gene expression levels by designing corresponding synthetic DNA sequences. Thanks to the recent community-wide effort [1], we are now able to elucidate the associations between yeast promoter sequences and the corresponding expression levels with the help of advanced deep learning methods. This milestone allows us to further explore the possibility of designing synthetic functional promoter sequences.

The conventional synthetic promoter sequence generation methods are based on pre-defined libraries of known promoters. Redden and Alper (2015) [2] assembled synthetic minimal core promoters using elements that were derived from different libraries. Rajkumar et al. (2016) [3] designed synthetic yeast promoters that are responsive to environmental stress according to known transcription factor binding-associated motifs information. Portela et al. (2017) [4] also designed synthetic promoters by analysing known promoters in libraries. All those conventional methods require task-specific library selection and complex example sequence analysis, on top of the limitations on synthetic sequence generation efficiency. More recently, with the help of deep learning methods, the efficiency and capacity of synthetic promoter design methods have been improved. For example, Kotopka and Smolke (2020) [5] trained an ensemble of convolutional neural networks by using 1 million yeast promoter sequences to design synthetic promoter sequences. However, due to the bottleneck of the sequence-to-expression predictive accuracy (i.e. an average squared Pearson correlation coefficient *r*^2^ value of 0.81), the performance and capacity of the predictive model-guided synthetic promoter design system still needs to be improved. Zrimec et al. (2022) [6] used generative adversarial networks which were also guided by a sequence-to-expression predictive model to generate synthetic yeast promoter sequences. However, analogously, the performance of the sequence-to-expression predictive model still needed to be further improved, due to a squared Pearson correlation coefficient *r*^2^ value of 0.80, leading to a bottleneck of the corresponding generative model’s performance. In addition, due to the natural difficulty on GAN training, the computational cost for obtaining the well-trained generative model was very high – the maximum value of the training iterations reached 200,000. More recently, two new GAN-based promoter generation methods namely DeepSEED [7] and PromoDGDE [8] were also proposed. The former successfully generated diverse and functional *Escherichia coli* promoter sequences, whilst the latter further extended the pipeline of the former to successfully generate yeast synthetic promoter sequences. However, both DeepSEED and PromoDGDE request extremely high computational cost as they rely on a combination of multiple AI methods. For example, PromoDGDE integrates GAN and Diffusion model with using a reinforce learning framework and an evolutionary algorithm-based optimization method. Analogously, DaSilva et al. (2025) [9] also proposed a diffusion model-based DNA sequence design method requiring high computational cost, due to its complex model architecture consisting of multiple ResNet blocks and attention layers, in addition to the standard diffusion model learning processes.

Temporal Convolutional Networks (TCN) [10] is a variant of convolutional neural networks for coping with sequence data. It demonstrates better performance over many popular sequence-based models on a variety of sequence data modelling tasks [11], with a simpler architecture and better computational efficiency. In general, TCN is constructed as a type of stacked neural networks, where each hidden layer has the same length as the input layer in order to guarantee that the prediction for the target time point *τ_i_* only depends on the information of all previous time-points (i.e. *τ*_0_, *τ*_1_, *τ*_2_, …, *τ_i−_*_1_). This type of training paradigm allows TCN to model the probabilistic distributions of all time-points of full sequences via the well-known autoregressive learning approach. TCN has already been used for different sequence data generative tasks [10] including speech generation, text-to-speech generation and music generation. In terms of handling biological sequence data, Shin, et al. (2021) [12] successfully adopted a TCN-based autoregressive model to estimate protein amino acid distributions and generate synthetic protein sequences.

In this work, we present a new TCN-based autoregressive generative model, namely Gen-DNA-TCN, which exploits a type of functional promoter sequence information (i.e. sequence-to-expression associations) that is encoded by Pre-DNA-TCN – an autoregressive predictive model that was developed for the recent Random Promoter DREAM Challenge [1]. The Pre-DNA-TCN model consists of 4.1 million parameters and well approximates the expression levels of random promoter sequences, according to a squared Pearson correlation coefficient *r*^2^ value of 0.88 and a Spearman correlation coefficient *ρ* value of 0.94 obtained over 71,103 high-quality testing yeast promoter sequences. By using the parameters of the pre-trained Pre-DNA-TCN model, we adopt a type of generative architecture and successfully conduct further training to obtain the Gen-DNA-TCN model with a very low computational cost. A large-scale evaluation confirms that Gen-DNA-TCN successfully generates a large number of unique, diverse and functional synthetic yeast promoter sequences that also encode similar transcription factor binding site distributions compared with real yeast promoter sequences.

## 2 Methods

### 2.1 Overview of Gen-DNA-TCN

In general, as shown in Figure 1.A, the proposed Gen-DNA-TCN model is a type of autoregressive generative network that is built on top of a pre-trained autoregressive predictive model, namely Pre-DNA-TCN [13], which successfully captured the associations between regulatory units of yeast promoter sequences and their corresponding expression levels. Gen-DNA-TCN exploits the 4.1 million parameters of the Pre-DNA-TCN model as prior knowledge to conduct further training by using 56,879 real yeast promoter sequences as the training dataset and 14,224 real yeast promoter sequences as the validation dataset for selecting the optimal model.

**Fig. 1:**
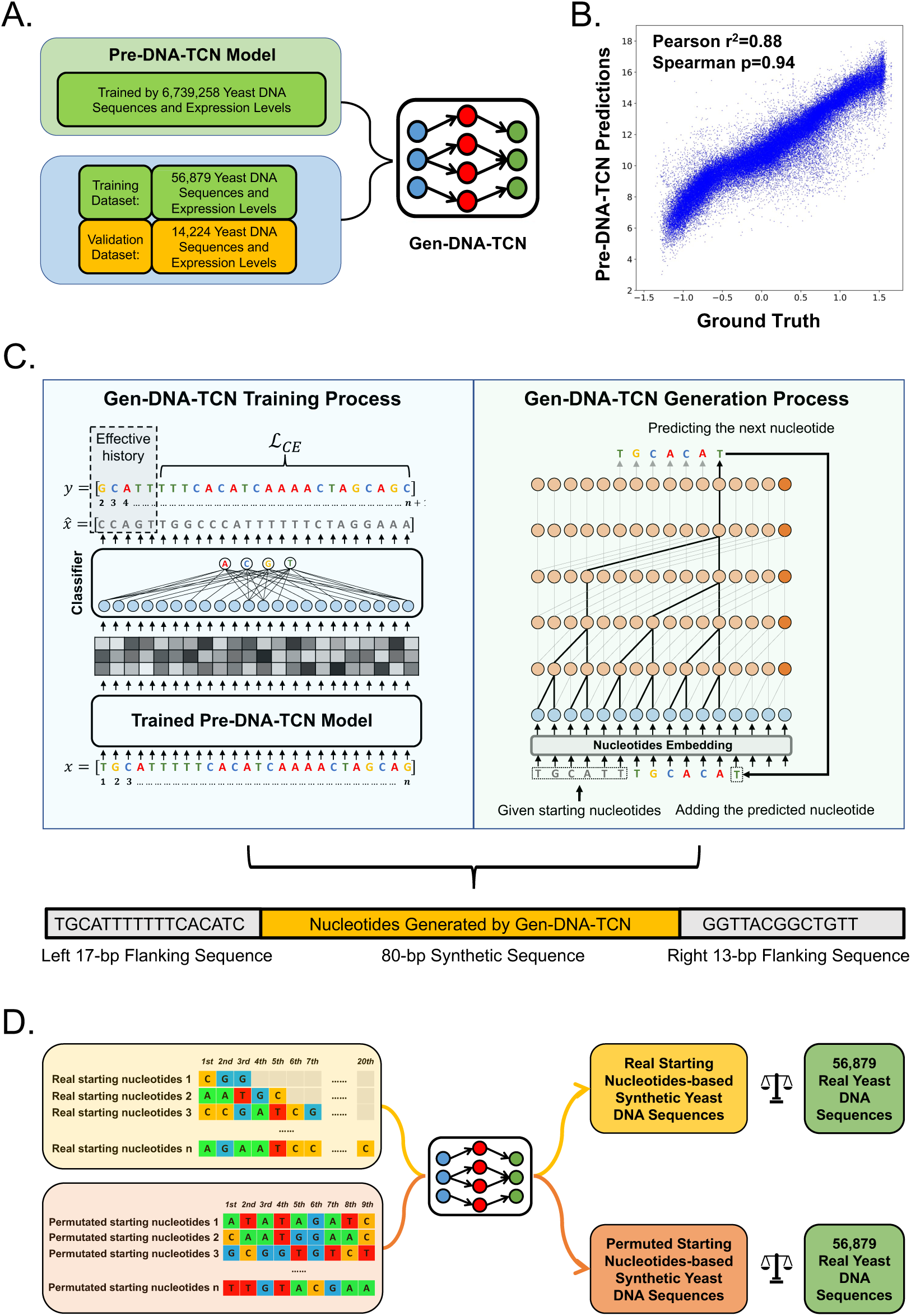
(A) The overview of Gen-DNA-TCN. (B) The predictive performance of the pre-trained Pre-DNA-TCN model on the 71,103 testing yeast promoter sequences. (C) The architecture of Gen-DNA-TCN and its training and sequence generation processes. (D) The performance evaluation approach for Gen-DNA-TCN.

### 2.2 The Pre-DNA-TCN model

The Pre-DNA-TCN model [13] is an autoregressive predictive model that encodes the associations between DNA sequences and their corresponding expression levels. It adopts a modified character-level temporal convolutional networks architecture and was trained by using 6.7 million random yeast promoter sequences along with their corresponding expression levels. In the recent Random Promoter DREAM Challenge [1] – a community-wide evaluation of deep learning-based sequence-to-expression predictive models, Pre-DNA-TCN obtained a squared Pearson score *r*^2^ of 0.88 and a Spearman score *ρ* of 0.94, demonstrating good performance in predicting the expression levels of 71,103 testing yeast promoter sequences. As shown in Figure 1.B, the predicted gene expression levels of the 71,103 testing yeast promoter sequences by Pre-DNA-TCN are very similar to their real expression levels. Such high predictive accuracy indicates the fact that the parameters *θ_pre_*of Pre-DNA-TCN successfully encode functional information about yeast promoter sequences.

### 2.3 The Gen-DNA-TCN model

Gen-DNA-TCN is a type of generative temporal convolutional networks that conduct autoregressive learning by directly exploiting the embedding layers of the pre-trained Pre-DNA-TCN model. As shown in Equation 1, let *x*=*{x*_1_, *x*_2_, …, *x_n_}* be a DNA sequence of length *n*, Gen-DNA-TCN learns the probability of the *i^th^* nucleotide considering all previous *i*-1 nucleotides’ distribution. *θ_Pre_ → θ_Gen_*denotes that the finally obtained parameters of Gen-DNA-TCN are derived from Pre-DNA-TCN’s parameters *θ_Pre_*. As *θ_Pre_* has already encoded the associations between promoter sequences and their corresponding expression levels, *θ_Gen_* actually encodes the probability distributions of the functional nucleotide combinations of yeast promoter sequences. Figure 1.C shows the architecture of the Gen-DNA-TCN model. In terms of its training process, Gen-DNA-TCN first takes *x* into the pre-trained Pre-DNA-TCN model to obtain a *d*-dimensional feature representation vector for each nucleotide in *x*, where *d* equals 384 in this work. The feature representation vector is further fed into a multi-class classification head, which consists of a *d*-dimensional linear layer and an output layer with four nodes representing four different nucleotides A, C, G, and T, respectively. The output logits are normalised using a Softmax function transforming them into probabilities. We model the target sequence *y* to be generated by the input sequence *x* but shifted by one nucleotide position, i.e. *y*=*{y*_2_, *y*_3_, …, *y_n_*_+1_*}*. Each nucleotide *y_i_* in the target sequence is predicted given all previous nucleotides. The well-known cross-entropy *L_CE_* is used as the loss function considering *y* and *x̂*, which denotes the output of the model. We define the first five initial nucleotides as effective history which is required to guarantee causality, so that the nucleotides generation process can be trained starting from the 6*^th^* nucleotide, and the effective history does not contribute to the loss function value. The model is trained by 1,000 epochs using the Adam optimiser with a learning rate of 0.002. We use 71,103 random promoter DNA sequences (each with a length of 110 nucleotides) to train the Gen-DNA-TCN model. Those sequences were also used as the testing dataset in the Random Promoter DREAM Challenge [1] and were not used to train the Pre-DNA-TCN model. We further split all 71,103 sequences into a training dataset and a validation dataset with a proportion of 8:2. We then preprocessed those 56,879 training sequences and 14,224 validation sequences by removing the left flanking sequences consisting of 17 nucleotides (i.e. TGCATTTTTTTCACATC), leading to two new sets of DNA sequences all consisting of 93 nucleotides. Note that, we still keep the right flanking sequences consisting of 13 nucleotides (i.e. GGTTACGGCTGTT) during the training process as a type of constraint. After every training epoch, we freeze the Gen-DNA-TCN parameters and compute the cross entropy value using the validation sequences. After completing 1,000 training epochs, we select the optimal model that obtains the lowest cross entropy value.

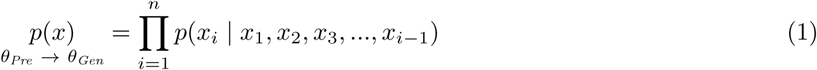

In terms of the Gen-DNA-TCN generation process, we adopt a type of sequential input-to-output mechanism to obtain synthetic sequences all consisting of 80 nucleotides, i.e. 110 bp nucleotides exclude the left 17 nucleotides and the right 13 nucleotides that are used as flanking sequences. Therefore, given a synthetic DNA sequence with a length of 80, we define two segments, i.e. a segment consisting of a set of pre-defined *n* starting nucleotides and another segment consisting of a set of corresponding 80*−n* generated nucleotides. We first use those pre-defined *n* starting nucleotides to generate the (*n*+1)*^th^* nucleotide, which is then used to form a new set of starting nucleotides consisting of *n*+1 nucleotides to generate the (*n*+2)*^th^* nucleotide. This process continues until reaching a pre-defined length *n_s_* of the generated sequences. In this paper, we define the value of *n_s_* equals 93. Finally, the left flanking sequences are appended in the beginning of the synthetic sequences, whilst the rightmost 13 generated nucleotides are replaced by the right flanking sequences, leading to 110 nucleotides in total, which are consistent with the settings of the Random Promoter DREAM Challenge [1]. For example, as shown in Figure 1.C, given a set of 6 starting nucleotides as inputs (i.e. TGCATT), Gen-DNA-TCN first generates the 7*^th^* nucleotide (i.e. T), which is appended with those 6 previous nucleotides as new inputs to generate the 8*^th^* nucleotide (i.e. G). The process terminates when the 93*^rd^* nucleotide of the synthetic sequence is generated.

### 2.4 Computational Experiments

We evaluate the performance of the Gen-DNA-TCN model by training two more variants of the proposed Gen-DNA-TCN models using different training datasets. As shown in Table 1, the original Gen-DNA-TCN model is denoted as Model 1, whilst Model 2 denotes a model that exploits the 56,879 sequences for both Pre-DNA-TCN model pre-training and for the second stage of the Gen-DNA-TCN model training. In addition, Model 3 does not use the pre-trained Pre-DNA-TCN model and directly exploits the 56,879 sequences to train the Gen-DNA-TCN model. In terms of the information encoded in Model 3, its parameters are randomly initialised at the beginning of the training process. Therefore, those parameters do not encode any functional information whilst merely capture the probability distributions of nucleotide combinations.

**Table 1:**
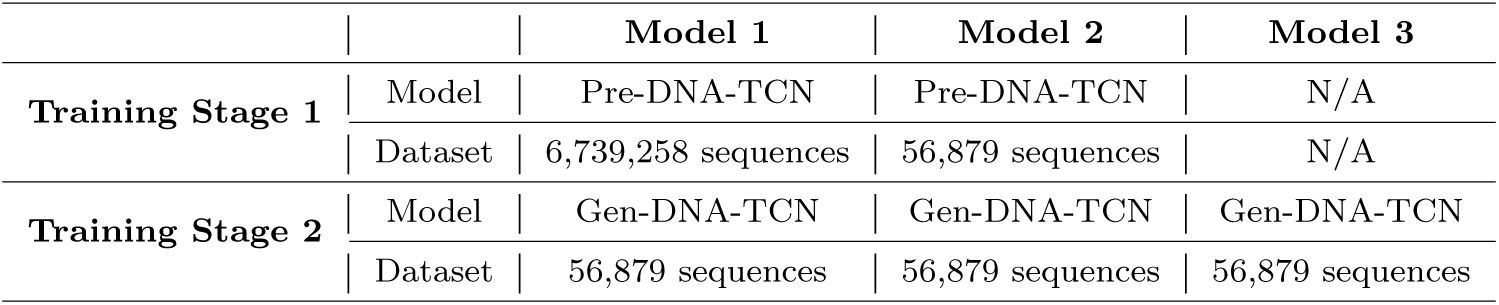
The training datasets used by different variants of Gen-DNA-TCN.

We also adopt two types of starting nucleotide combinations to generate synthetic sequences. As shown in Figure 1.D, the first type of starting nucleotides are extracted from the real 56,879 training sequences, whilst the second type of starting nucleotides are made by random permutations. As shown in Table 2, the numbers of unique starting nucleotides that are derived from real sequences vary according to different starting nucleotide combinations. For example, there exist 64 unique combinations of 3 starting nucleotides, but the number of unique combinations of 20 starting nucleotides reaches to 24,154. We generate synthetic sequences by using the real starting nucleotides whose lengths range from 1 to 20, and remove any duplicated sequences w.r.t. both the real and synthetic sequences.

**Table 2:**
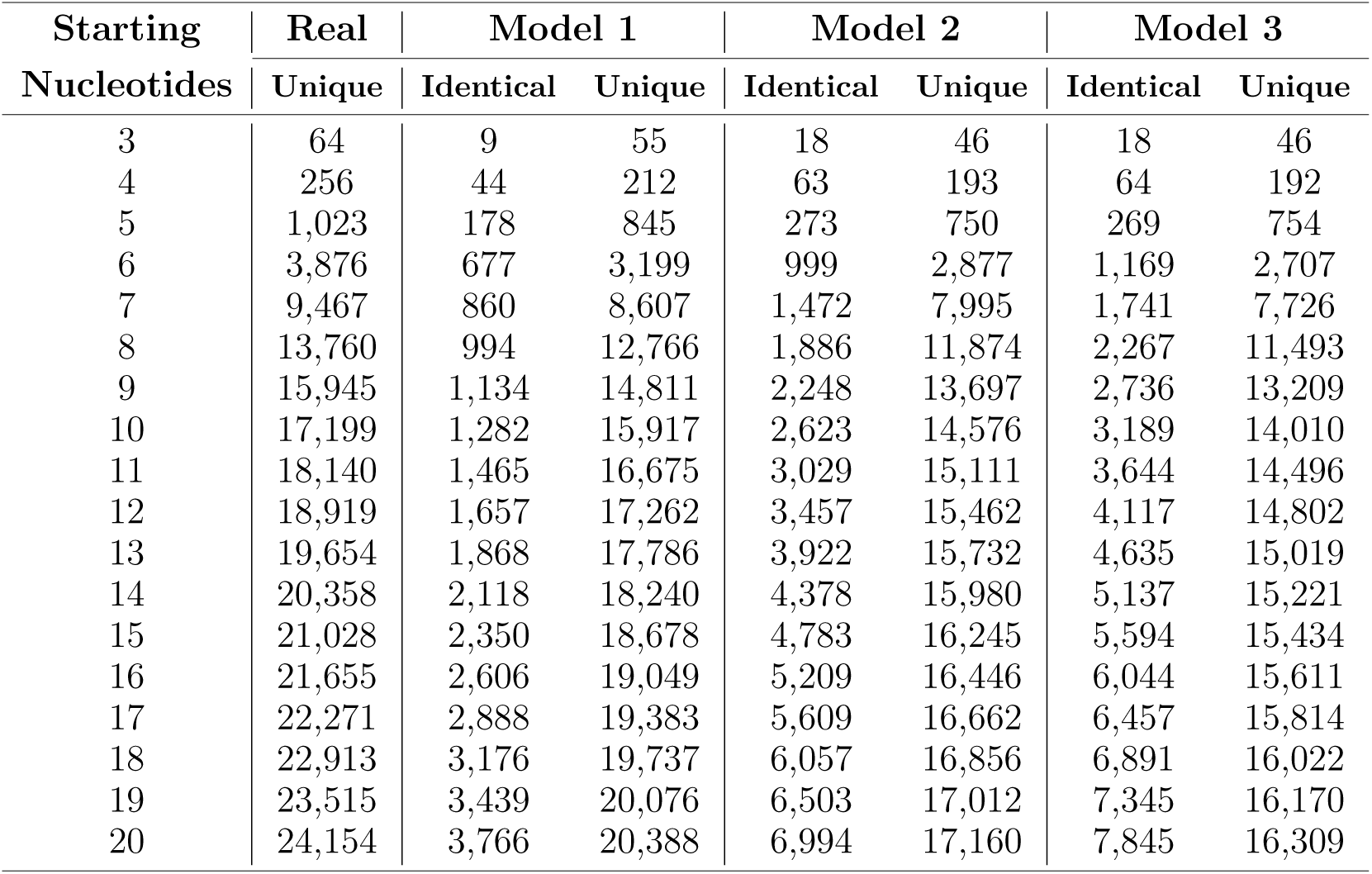
The number of identical and unique synthetic promoter sequences generated by three models using different real starting nucleotide combinations.

The second type of starting nucleotides is made by randomly permutating the four different nucleotides. We generate 262,144 synthetic sequences using the combinations of 9 starting nucleotides, since the total number of starting nucleotide combinations equals to *k^h^*, where *k* is the number of different nucleotides and *h* is the length of the starting sequence, leading to 4^9^ (i.e. 262,144) combinations, which are more than the number of the total real promoter sequences (i.e. 71,103). However, if *k <* 9, the number of starting nucleotide combinations will be less than the number of the total real promoter sequences. We then discard the left-most 9 random permutated starting nucleotides and replace the right-most nucleotides with the right flanking sequences, after appending the left flanking sequences in the beginning, making the length of all synthetic sequences equal 110. Finally, we filter out any duplicated synthetic sequences w.r.t. the 71,103 training sequences for Gen-DNA-TCN and the 6.7 million training sequences for the pre-trained Pre-DNA-TCN model

We evaluate the quality of the generated synthetic sequences from three different aspects. To begin with, we compare the expression levels of the generated synthetic sequences with the expression levels of the real sequences. We use the pre-trained Pre-DNA-TCN model to predict the expression levels of the synthetic sequences, due to its high predictive accuracy. As the real sequences’ expression levels are in different scales, we also use Pre-DNA-TCN to obtain the expression levels of the real sequences for the comparison purpose. In order to facilitate the discussions, we create 11 bins of expression levels according to the range of the Pre-DNA-TCN-predicted expression levels for both real and synthetic sequences, i.e. 3.03 - 19.56. Moreover, we evaluate the distributions of transcription factor binding sites that are included in both the synthetic and real sequences. We search the transcription factor binding site motifs encoded in the real and synthetic sequences using the well-known JASPER yeast transcription factor binding site database (the 2024 version) [14] with FIMO [15]. We use a conservative adjusted significance level, i.e. *α*=5.00E-06, to confirm the detected transcription factor binding site motifs, due to the large number of the target sequences and the well-known multiple comparison problem in statistical tests. Furthermore, we evaluate the differences between the synthetic and real sequences by calculating the well-known Levenshtein distance [16] for all possible pairs of the synthetic and real sequences. As all the real and synthetic sequences have the same length of 110, the Levenshtein distance between any pair of sequences means the number of positions where different nucleotides are included in two sequences.

## 3 Results

### 3.1 Gen-DNA-TCN successfully generates functional synthetic sequences using real starting nucleotide combinations

In general, Gen-DNA-TCN successfully learns the distributions of yeast promoter sequences’ transcription factor binding sites, and is able to generate high-quality synthetic promoter sequences using real starting nucleotide combinations. To begin with, we first evaluate the quality of the synthetic sequences by comparing their expression profiles with the real sequences’ expression profiles. Figures 2.A - 2.C show the comparisons of the distributions of the Pre-DNA-TCN-predicted expression levels between the real sequences and the synthetic sequences that are generated by three different models. Table 3 also shows the actual numbers of the real and synthetic sequences that are predicted to have certain expression levels belonging to different bins, and the rankings of different bins according to the corresponding quantities of sequences, i.e. the bin including the fewest sequences is ranked as the 1*^st^*, whilst the bin including the most sequences is ranked as the 10*^th^*. Note that, the real sequences do not have their expression profiles being within the range of bin 11, and Model 1 also merely generated 4 synthetic sequences that belong to bin 11. On the contrary, both Model 2 and Model 3 generated more synthetic sequences belonging to bin 11, i.e. 40 and 48 respectively, showing more differences to the real sequences. Therefore, hereafter, we only discuss the synthetic sequences that belong to the first 10 bins. It is obvious that Model 1 generated the most unique synthetic sequences (i.e. 107,711 synthetic sequences in total excluding 4 sequences in bin 11), which also show the most similar distribution to the expression levels of the 56,879 real sequences. However, the synthetic sequences generated by either Model 2 or Model 3 show very different distributions, compared with the real sequences’ expression levels. It is also obvious that the Model 1-generated synthetic sequences have the same rankings with the real sequences in six bins, i.e. bins 1, 2, 4, 5, 6 and 10. Both the real and Model 1-generated synthetic sequences have the most and the second-most sequences distributed in bins 5 and 6, respectively. However, the synthetic sequences generated by Model 2 and Model 3 respectively only have the same rankings with the real sequences in four bins, i.e. bins 1, 3, 4 and 9. There exist the most and the second-most synthetic sequences generated by Model 2 and Model 3 respectively in bins 8 and 7, rather than in bins 5 and 6. This patterns suggest that the expression profiles of the synthetic sequences generated by both Model 2 and Model 3 follow very different distributions to the expression profiles of the real sequences.

**Fig. 2:**
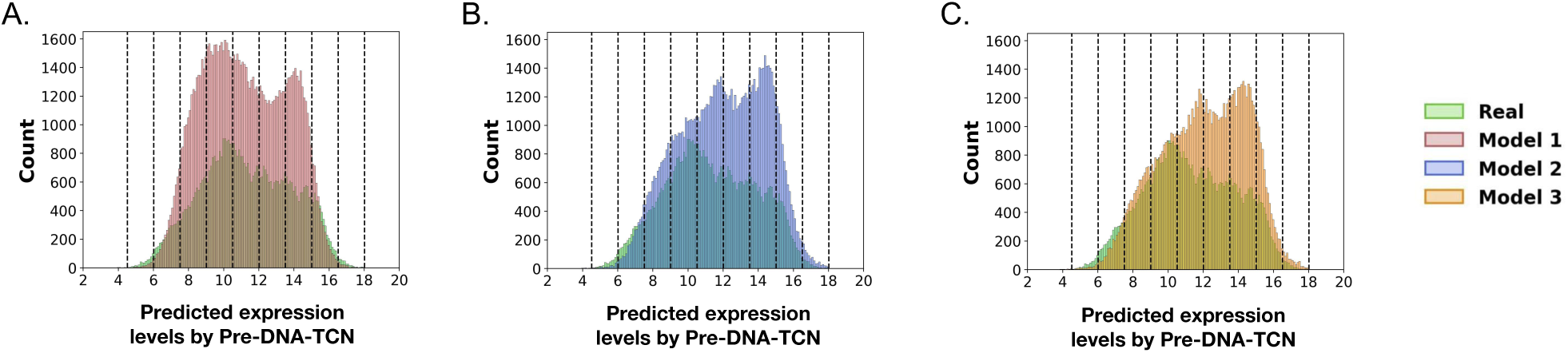
The predicted expression levels of the real and the synthetic sequences generated by Model 1 (A), Model 2 (B) and Model 3 (C) using the real starting nucleotides.

**Table 3:** The numbers of unique synthetic promoter sequences generated by three models using the real starting nucleotides according to the pre-defined bins of expression levels.

| Bins | Expression Level | Real Sequences |  | Model 1<br>Synthetic Sequences |  | Model 2<br>Synthetic Sequences |  | Model 3<br>Synthetic Sequences |  |
| --- | --- | --- | --- | --- | --- | --- | --- | --- | --- |
|  | Range | Quantity | Ranking | Quantity | Ranking | Quantity | Ranking | Quantity | Ranking |
| 1 | 3.03 - 4.52 | 17 | 1 | 7 | 1 | 13 | 1 | 21 | 1 |
| 2 | 4.52 - 6.02 | 521 | 3 | 231 | 3 | 205 | 2 | 241 | 2 |
| 3 | 6.02 - 7.52 | 3,536 | 4 | 4,740 | 5 | 3,118 | 4 | 2,703 | 4 |
| 4 | 7.52 - 9.02 | 7,438 | 6 | 17,775 | 6 | 10,071 | 6 | 8,563 | 6 |
| 5 | 9.02 - 10.52 | 11,928 | 10 | 23,009 | 10 | 14,725 | 7 | 12,681 | 7 |
| 6 | 10.52 - 12.02 | 11,003 | 9 | 20,501 | 9 | 17,872 | 8 | 15,975 | 8 |
| 7 | 12.02 - 13.52 | 9,327 | 8 | 17,956 | 7 | 17,905 | 9 | 16,526 | 9 |
| 8 | 13.52 - 15.02 | 8,281 | 7 | 18,662 | 8 | 20,112 | 10 | 18,453 | 10 |
| 9 | 15.02 - 16.52 | 4,488 | 5 | 4,668 | 4 | 8,864 | 5 | 8,002 | 5 |
| 10 | 16.52 - 18.02 | 340 | 2 | 162 | 2 | 929 | 3 | 839 | 3 |
| * 11 | 18.02 - 19.56 | 0 | N/A | 4 | N/A | 40 | N/A | 48 | N/A |
| Total | 3.03 - 19.56 | 56,879 | N/A | 107,715 | N/A | 93,854 | N/A | 84,052 | N/A |

Moreover, we further investigate the distributions of transcription factor binding sites considering their position information over the synthetic DNA sequences that all consist of 110 nucleotides. As shown in Figure 3.A, in terms of the positive strand results, both the red and the green distributions are very similar, suggesting that Model 1 accurately learns the transcription factor binding site position distributions over the entire 110 nucleotides, except some areas such as between the 30*^th^* and the 40*^th^*bp, and between the 55*^th^* and the 60*^th^*bp. Analogously, in terms of the negative strand results, both the red and green distributions are very similar, as shown in Figure 3.B. On the contrary, the synthetic sequences generated by both Model 2 and Model 3 include very different transcription factor binding site position distributions to the real sequences. As shown in Figures 3.C - 3.F, the blue and orange distributions are very different to the green distribution, suggesting both Model 2 and Model 3 fail to learn the position distributions of the transcription factor binding sites included in the real sequences.

**Fig. 3:**
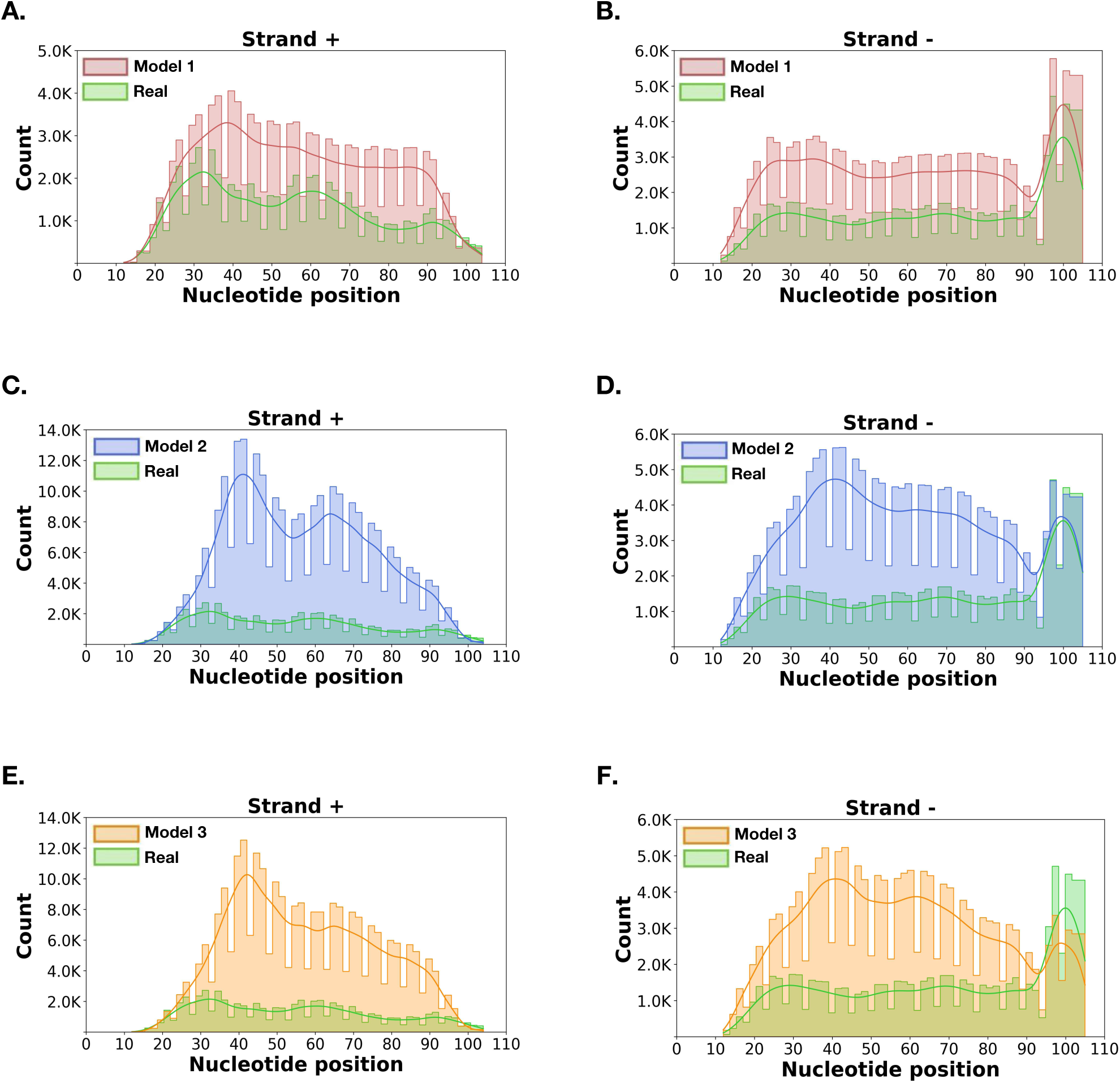
The position distributions of the transcription factor binding sites detected in positive and negative strands of the real sequences and the synthetic sequences generated by Model 1 (A-B), Model 2 (C-D) and Model 3 (E-F) using the real starting nucleotides.

Furthermore, we investigate the details of the transcription factor binding sites that are included in the synthetic and real sequences. In general, the real sequences include 50 and 51 different transcription factor binding site motifs, respectively according to the positive and negative strands results using the conservative significance level, i.e. *α*=5.00E-06. The synthetic sequences generated by Model 1, Model 2 and Model 3 include almost the same types of transcription factor binding site motifs as the real sequences, except motif MGA1, which is not detected in the real sequences. In addition, as different models generate different numbers of synthetic sequences, we normalise the exact frequencies of different motifs by dividing the total number of sequences. For example, the exact frequency of motif *HAP5* detected in the positive strands of the 107,711 Model 1-generated synthetic sequences equals 95. Hence, the normalised frequency value approximately equals 8.82 E-04.

Among those three models, the transcription factor binding sites detected in the positive strands of Model 1’s synthetic sequences show the strongest associations with the transcription factor binding sites included in the real sequences as denoted by the highest Pearson correlation coefficient value (i.e. *r*=0.818), whereas Model 2 and Model 3 both show weaker associations due to the lower Pearson correlation coefficient values, i.e. *r*=0.721 and *r*=0.766. Among Figures 4.A - 4.C, the distributions of the dots in Figure 4.A are the most compact, suggesting the most similar distributions of the detected transcription factor binding sites in Model 1-generated synthetic sequences and the real sequences. In addition, Figure 4.G shows the comparisons of those high-frequent transcription factor binding sites. It is obvious that Model 1 shows the best performance, whereas Model 2 and Model 3 tend to generate many more synthetic sequences including certain motifs such as *REB1* and *NSI1*, as denoted by the dramatically higher bars.

**Fig. 4:**
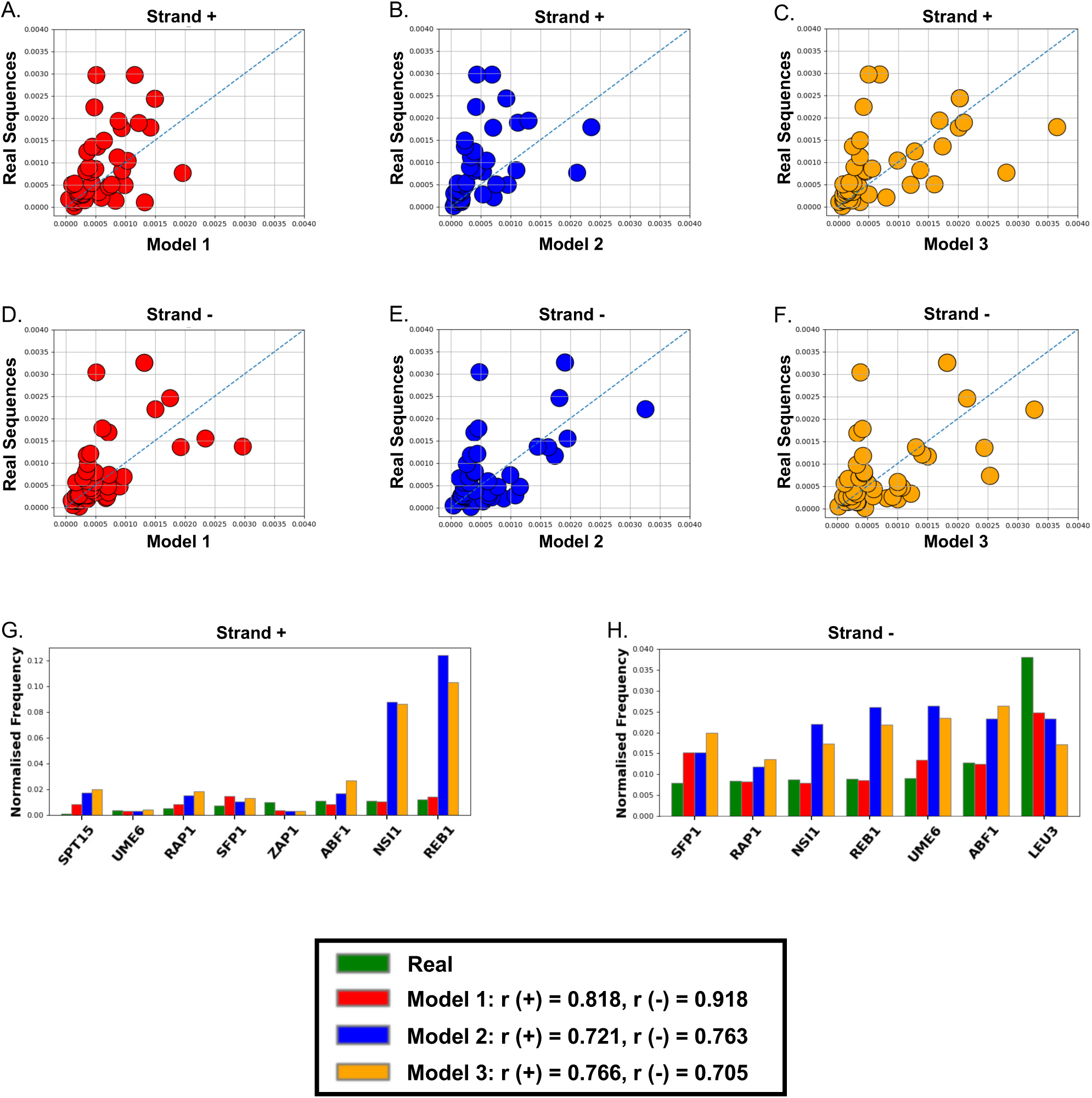
The pairwise comparisons of the normalised frequencies of the transcription factor binding sites detected in the positive strands (A-C) and the negative strands (D-F) of the real and synthetic sequences generated by Model 1, Model 2 and Model 3 using the real starting nucleotides. The comparisons of the normalised frequencies of certain high-frequent transcription factor binding sites detected in the positive strands (G) and the negative strands (H) of the real and synthetic sequences generated by Model 1, Model 2 and Model 3 using the real starting nucleotides.

In terms of the negative strand results, the transcription factor binding sites included in the synthetic sequences generated by Model 1 also show the strongest associations with the real sequences due to the highest Pearson correlation coefficient value (i.e. *r*=0.918), whereas Model 2 and Model 3 both show lower Pearson correlation coefficient values, i.e. *r*=0.763 and *r*=0.705. Analogously, the distributions of the dots in Figure 4.D are the most compact, compared with Figures 4.E and 4.F. In terms of those high-frequent motifs, as shown in Figure 4.H, Model 1 also shows the best performance, since the distributions of those motifs are the most similar to the distributions of the corresponding motifs included in the real sequences. For example, the frequencies of motifs *ABF1*, *UME6*, *REB1* and *NSI1* included in the synthetic sequences generated by Model 2 and Model 3 are all much higher than the ones included in both the real sequences and the Model 1-generated synthetic sequences.

### 3.2 Gen-DNA-TCN successfully generates diverse functional synthetic sequences that are different to real sequences

As another crucial performance evaluation criterion for generative models, we further investigate the differences between the real sequences and the synthetic sequences generated by all three models. We calculate the Levenshtein distance [16] between all possible pairs of the real and synthetic sequences, i.e. 56,879 real sequences versus 107,711 synthetic sequences for Model 1, 56,879 real sequences versus 93,814 synthetic sequences for Model 2, and 56,879 real sequences versus 84,004 synthetic sequences for Model 3. Note that, as all sequences have the same left and right flanking sequences, the differences between sequences are considered only in the middle of 80 synthetic nucleotides.

In general, the synthetic sequences generated by the three different models all have high differences to the real sequences. As shown in Figures 5.A - 5.C, the embeddings of the *t*-SNE-transformed [17] real and synthetic sequences respectively generated by all three different models are distributed in different areas. In terms of the Levenshtein distance values, as shown in Figure 5.D, the median Levenshtein distance values for the synthetic sequences generated by three different models all equal 46.0, whilst the mean values equal 46.2, 46.5 and 46.5, respectively. Figure 5.E shows the boxplots regarding the Levenshtein distance values between the real and the synthetic sequences according to different bins of expression profiles. Overall, the synthetic sequences generated by all three models have the same median Levenshtein distance values ranging between 43.0 and 47.0 in six bins, i.e. bins 2, 3, 5, 7, 8 and 9. The Model 1-generated synthetic sequences have slightly higher median Levenshtein distance values than the synthetic sequences generated by Model 2 and Model 3 in two bins, i.e. 48.0 in bin 1 and 47.0 in bin 10. The synthetic sequences generated by Model 2 and Model 3 only have higher Levenshtein distance values than the Model 1-generated synthetic sequences in one bin, i.e. 47.0 in bin 6.

**Fig. 5:**
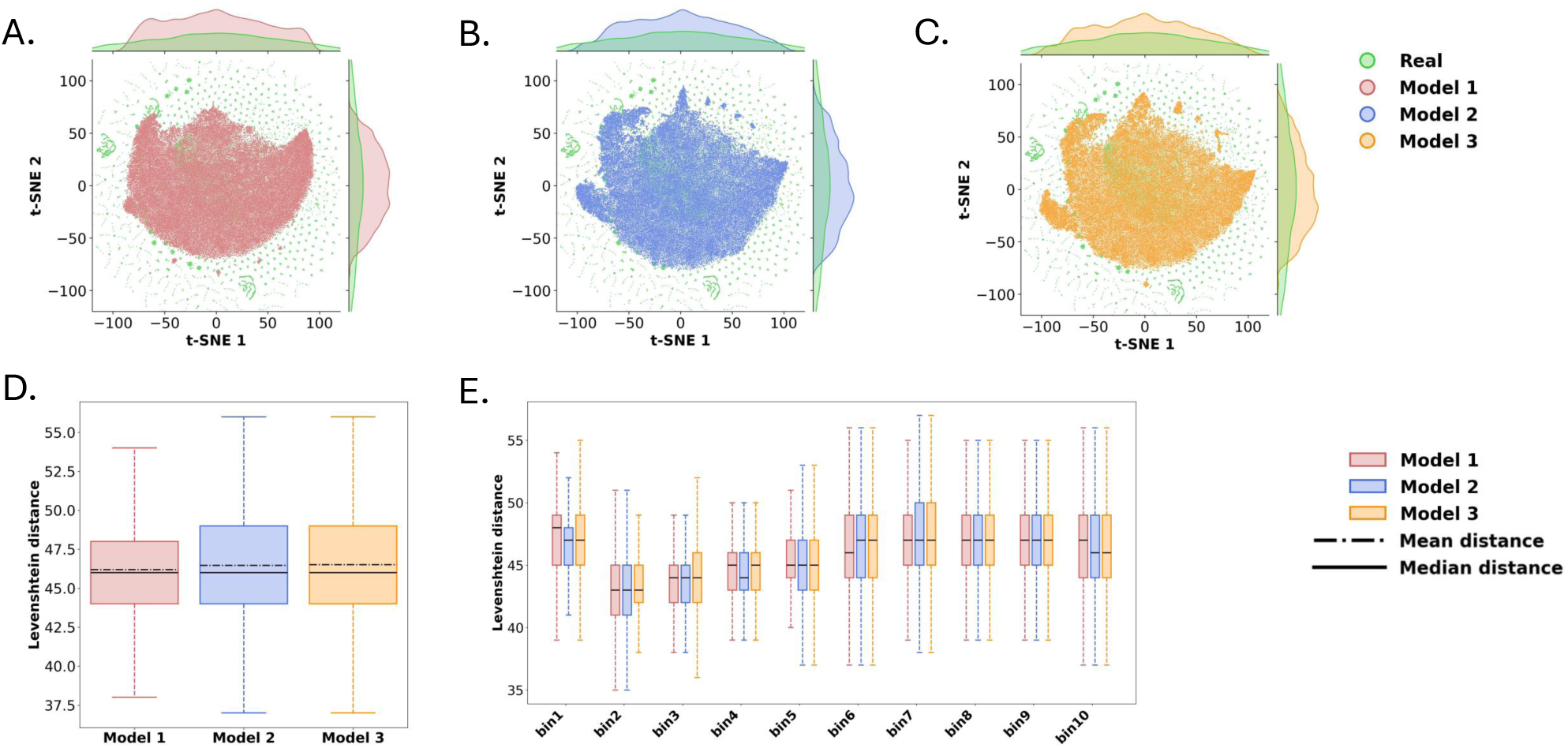
(A-C) The distributions of the *t*-SNE transformed embeddings of the real and synthetic sequences generated by Model 1, Model 2 and Model 3 using the real starting nucleotides. (D) The boxplots of the distributions of the Levenshtein distance values obtained by all possible pairs of real and synthetic sequences respectively generated by Model 1, Model 2 and Model 3 using the real starting nucleotides. (E) The expression level bin-based distributions of the Levenshtein distance values obtained by all possible pairs of real and synthetic sequences respectively generated by Model 1, Model 2 and Model 3 using the real starting nucleotides.

### 3.3 Gen-DNA-TCN shows good model generalisation ability and successfully generates functional synthetic sequences using randomly permutated starting nucleotide combinations

We then further evaluate the performance of Gen-DNA-TCN (Model 1) by using the randomly permutated starting nucleotide combinations as inputs to generate synthetic DNA sequences. As introduced in the Computational Experiments section, 262,144 synthetic sequences are initially generated, we further remove any duplicated sequences and those have expression levels being within bin 11, leading 261,487 unique synthetic sequences. We also create two subsets of the synthetic sequences by using two different sampling strategies, i.e. bin-based sampling and random sampling. In terms of the bin-based sampling strategy, we allocate the synthetic sequences into the different bins by considering their expression levels. For each bin that includes more synthetic sequences than the corresponding real sequences, we randomly select the same amount of the synthetic sequences. If a bin includes more real sequences than synthetic sequences, no sampling will be conducted for that bin. As shown in Table 4, 53,302 synthetic sequences are selected by using the bin-based sampling strategy. In terms of the random sampling strategy, 56,879 synthetic sequences are randomly picked from the entire synthetic sequence set.

**Table 4:** The numbers of unique synthetic promoter sequences generated by Model 1 using the permutated starting nucleotide combinations according to the pre-defined bins of expression levels.

| Bins | Expression Level | Real Sequences |  | All Generated Synthetic Sequences |  | Bin-based Sampled Synthetic Sequences |  | Randomly Sampled Synthetic Sequences |  |
| --- | --- | --- | --- | --- | --- | --- | --- | --- | --- |
|  | Range | Quantity | Ranking | Quantity | Ranking | Quantity | Ranking | Quantity | Ranking |
| 1 | 3.03 - 4.52 | 17 | 1 | 10 | 1 | 10 | 1 | 3 | 2 |
| 2 | 4.52 - 6.02 | 521 | 3 | 507 | 3 | 507 | 3 | 109 | 3 |
| 3 | 6.02 - 7.52 | 3,536 | 4 | 17,428 | 5 | 3,536 | 5 | 3,877 | 5 |
| 4 | 7.52 - 9.02 | 7,438 | 6 | 59,294 | 9 | 7,438 | 6 | 12,998 | 9 |
| 5 | 9.02 - 10.52 | 11,928 | 10 | 68,329 | 10 | 11,928 | 10 | 15,029 | 10 |
| 6 | 10.52 - 12.02 | 11,003 | 9 | 55,565 | 8 | 11,003 | 9 | 11,785 | 8 |
| 7 | 12.02 - 13.52 | 9,327 | 8 | 40,879 | 7 | 9,327 | 8 | 8,815 | 7 |
| 8 | 13.52 - 15.02 | 8,281 | 7 | 18,203 | 6 | 8,281 | 7 | 3,993 | 6 |
| 9 | 15.02 - 16.52 | 4,488 | 5 | 1,256 | 4 | 1,256 | 4 | 268 | 4 |
| 10 | 16.52 - 18.02 | 340 | 2 | 16 | 2 | 16 | 2 | 2 | 1 |
| Total | 3.03 - 18.02 | 56,879 | N/A | 261,487 | N/A | 53,302 | N/A | 56,879 | N/A |

In general, Gen-DNA-TCN shows good performance and model generalisation ability, since it can generate similar transcription factor binding sites included in the real sequences even using randomly permutated starting nucleotide combinations. To begin with, as shown in Figure 6.A and Table 4, both the synthetic sequences and the real sequences have their peak values in bin 5. In terms of the rankings of different bins, the real sequences and the synthetic sequences have the same rankings in 4 bins, i.e. bins 1, 2, 5 and 10. In terms of the bin-based sampled synthetic sequences, as shown in Figure 6.B and Table 4, their expression level distributions are more similar to the real sequences’ expression level distribution, as both the red and the green distributions have large overlapping areas in bins 1, 2, 3, 4, 5, 6 and 7. Both sets of sequences also have the same rankings in all bins except bins 3 and 9. In terms of the randomly sampled synthetic sequences, as shown in Figure 6.C, both the red and the green distributions have the large area of overlapping in bins 5, 6, and 7. Both sets of sequences have the same rankings in bins 2 and 5, as shown in Table 4.

**Fig. 6:**
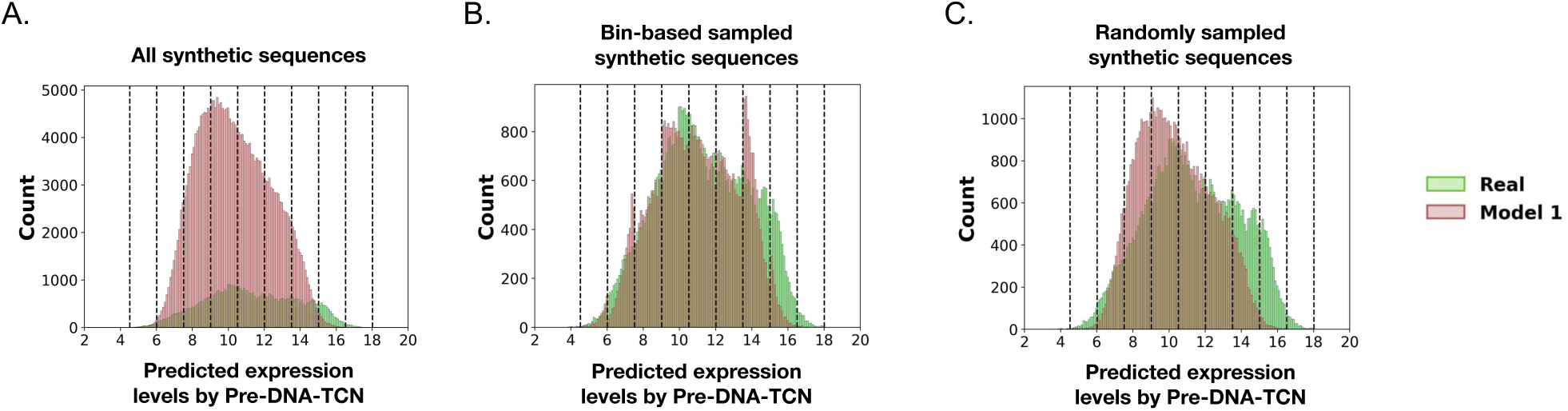
The predicted expression levels of the real and three groups of synthetic sequences generated by Model 1, i.e. the full generated synthetic sequences group (A), the bin-based sampled synthetic sequences group (B), and the randomly sampled synthetic sequences group (C), using the permutated starting nucleotides.

Moreover, in terms of the position distributions of the detected transcription factor binging sites, as shown in Figures 7.A and 7.B, both the real and the synthetic sequences show similar distributions, as denoted by the green and the red curves. In terms of both the positive and the negative strand results, both curves have different shapes between the 80*^th^* and the 110*^th^*bp. In terms of the bin-based sampled synthetic sequences, they have similar shapes to the real sequences in general. In terms of the positive strand results, as shown in Figure 7.C, both the red and the green distributions show similar patterns, but the synthetic sequences have more transcription factor binding sites than the real sequences in some areas. In terms of the negative strand results, as shown in Figure 7.D, both curves show similar patterns, but they have different shapes between the 90*^th^* and the 110*^th^* bp. Regarding the randomly sampled synthetic sequences, as shown in Figure 7.E, both curves show similar patterns, but the real and the synthetic sequences show very different shapes after the 50*^th^* bp. Analogously, in terms of the negative strand results, as shown in Figure 7.F, both the red and the green curves also show similar patterns, though the shapes of the two curves become very different between the 80*^th^*and the 110*^th^* bp.

**Fig. 7:**
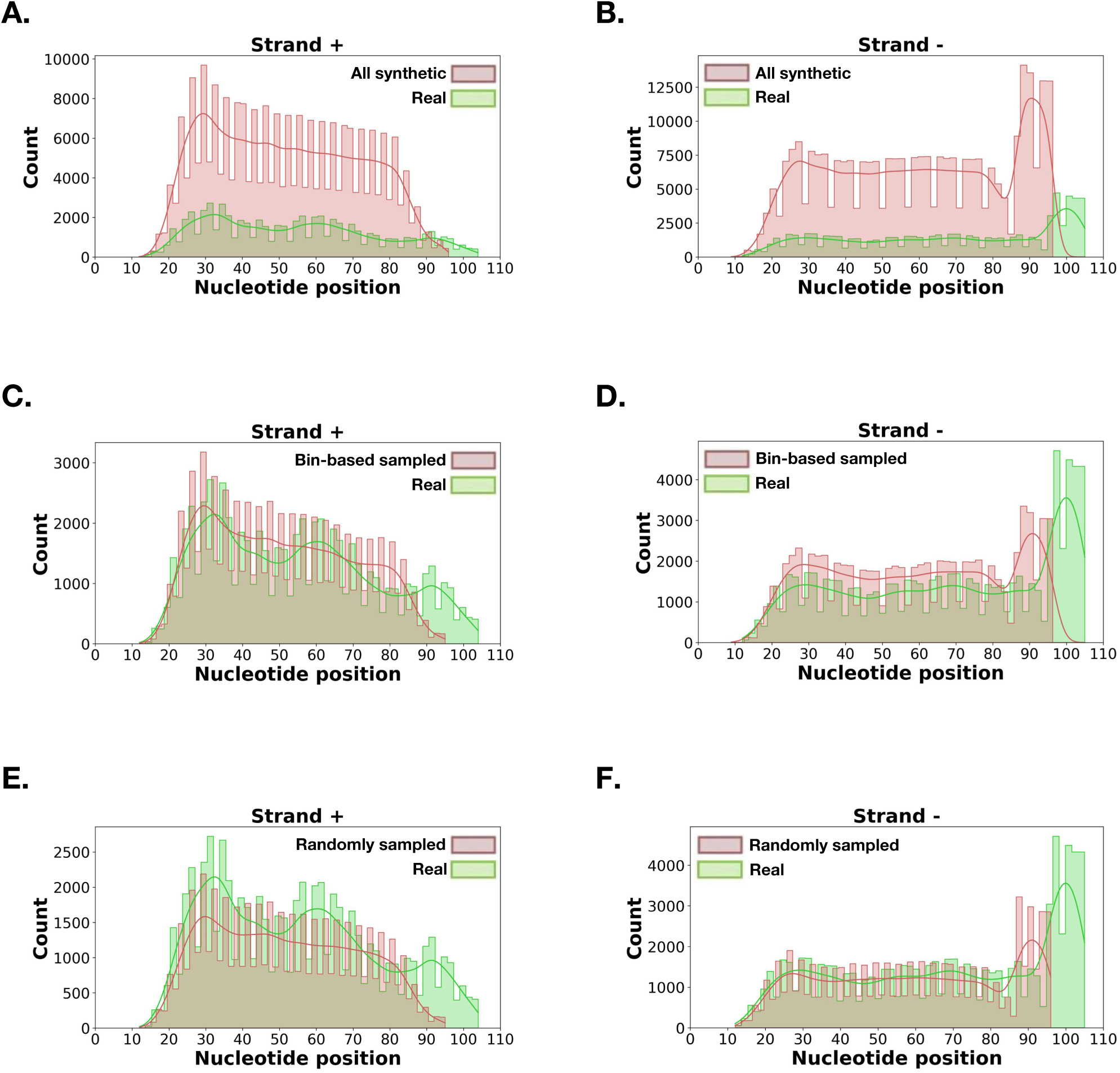
The position distributions of the transcription factor binding sites detected in positive and negative strands of the real sequences and the synthetic sequences in the full synthetic sequences group (A-B), the bin-based sampled synthetic sequences group (C-D) and the randomly sampled synthetic sequences group (E-F) generated using the permutated starting nucleotides.

Furthermore, in terms of the transcription factor binding sites detected by the JASPAR database and FIMO, the detected synthetic sequences in the three different groups have the same transcription factor binding site motifs as the real sequences, except motif MGA1 in the positive strands, which is not detected in the real sequences. All three groups of the synthetic sequences also show strong Pearson correlation coefficient values with the real sequences regarding the distributions of the detected transcription factor binding sites. As shown in Figures 8.A and 8.D, all the synthetic sequences show good correlations with the real sequences in both the positive and negative strands results, with the Pearson correlation coefficient values of 0.770 and 0.913, respectively. In terms of the synthetic sequences sampled by the bin-based strategy, as shown in Figures 8.B, they obtain the strongest Pearson correlation coefficient value (i.e. 0.829) for the positive strand results. However, they show a weaker Pearson correlation coefficient value (i.e. 0.894), as also shown in Figure 8.E. In terms of the synthetic sequences sampled by the random strategy, as shown in Figure 8.C, the Pearson correlation coefficient value for the positive strands, i.e. 0.762, is the lowest among all the three groups. However, the Pearson correlation coefficient value for the negative strands is the highest, i.e. 0.915.

**Fig. 8:**
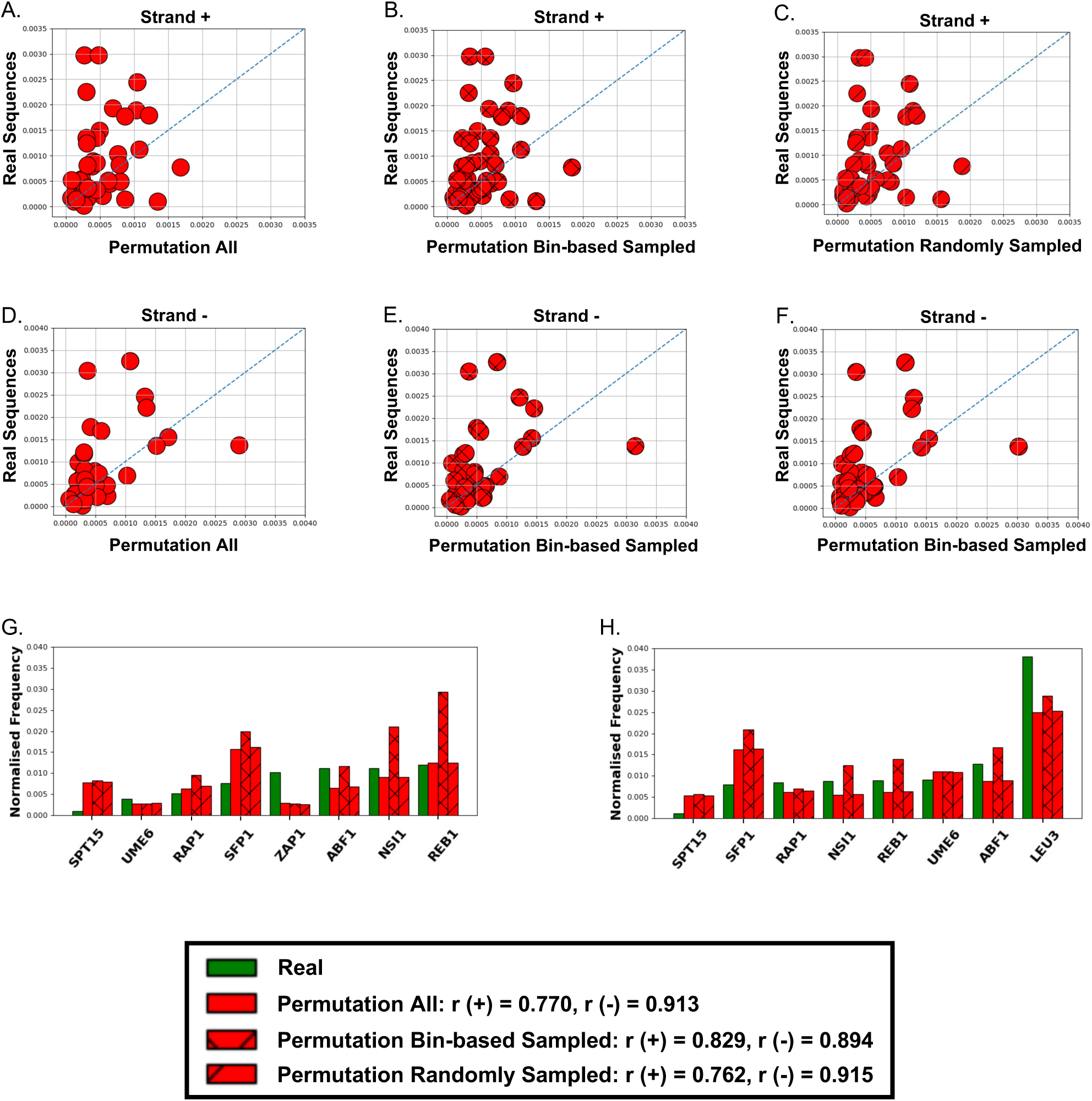
The pairwise comparisons of the normalised frequencies of the transcription factor binding sites detected in the positive strands (A-C) and the negative strands (D-F) of the real and three groups of synthetic sequences generated by Model 1 using the permutated starting nucleotides. The comparisons of the normalised frequencies of certain high-frequent transcription factor binding sites detected in the positive strands (G) and the negative strands (H) of the real and three groups of synthetic sequences generated by Model 1 using the permutated starting nucleotides.

### 3.4 The synthetic sequences generated by using the randomly permutated starting nucleotide combinations are also very different to the real sequences

We also compute the Levenshtein distance values between all possible pairs of the real and those three groups of synthetic sequences generated by the permutated starting nucleotide respectively, i.e. 56,879 real sequences versus 261,487 synthetic sequences, 56,879 real sequences versus 53,302 bin-based sampled synthetic sequences, and 56,879 real sequences versus 56,879 randomly sampled synthetic sequences.

In general, Gen-DNA-TCN also successfully generates diverse synthetic sequences using the randomly permutated starting nucleotide combinations. As shown in Figures 9.A - 9.C, the *t*-SNE-transformation-based 2D visualisations confirm that all three different groups of the synthetic sequences follow very different distributions to the real sequences. Figure 9.D shows the boxplots of the Levenshtein distance values of the pairwise comparisons. It is clear that the median Levenshtein distance values for all three groups of synthetic sequences all equal 46.0, whilst the mean Levenshtein distance values for all three groups of synthetic sequences are also very close to 46.0, i.e. 46.2, 46.4 and 46.2, respectively. In terms of the bin-based boxplots shown in Figure 9.E, the median Levenshtein distance values for all the synthetic sequences range between 43.0 and 48.0. The bin-based sampled synthetic sequences’ median Levenshtein distance values range between 44.0 and 48.0, whilst the median Levenshtein distance values for the randomly sampled synthetic sequences range between 43.0 and 47.0.

**Fig. 9:**
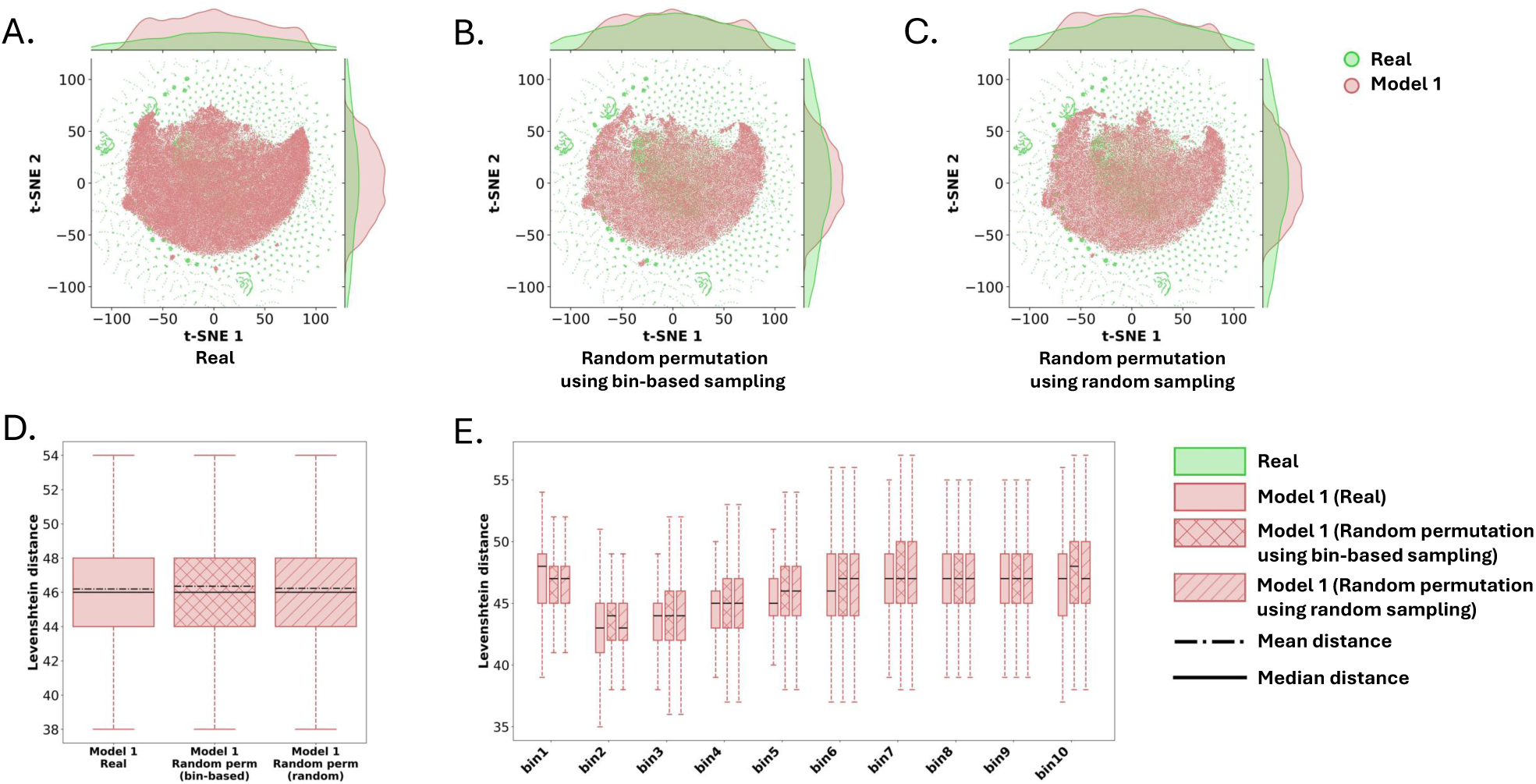
(A-C) The distributions of the t-SNE transformed embeddings of the real and three groups of synthetic sequences generated by Model 1 using the permutated starting nucleotides. (D) The boxplots of the distributions of the Levenshtein distance values obtained by all possible pairs of real and three different groups of synthetic sequences respectively generated by Model 1 using the permutated starting nucleotides. (E) The expression level bin-based distributions of the Levenshtein distance values obtained by all possible pairs of real and three different groups of synthetic sequences respectively generated by Model 1 using the permutated starting nucleotides.

## 4 Discussions

Overall, as introduced in previous sections, Gen-DNA-TCN successfully generated synthetic promoter sequences that bear similar distributions of transcription factor binding sites in real yeast promoter sequences. In this section, we further discuss the generated synthetic yeast promoter sequences and the computational efficiency of the Gen-DNA-TCN model training.

### 4.1 Gen-DNA-TCN shows the capability on generating synthetic promoter sequences bearing more than one transcription factor binding sites

Promoter sequences that include more than one transcription factor binding sites are usually related with more complex gene regulatory mechanisms. Gen-DNA-TCN is also able to generate synthetic promoter sequences including two transcription factor binding sites using either real starting nucleotides or randomly permunated starting nucleotides. Figure 10 shows four example synthetic promoter sequences that are generated by Gen-DNA-TCN. For example, when using the real starting nucleotides TTT, the generated sequence includes two transcription factor binding sites, i.e. *YRR1* and *SPT15*, in two separated parts of the positive strand, as highlighted. Analogously, when using the real starting nucleotides GGA, the generated sequence includes another two transcription factor binding sites, i.e. *STB3* and *ABF1*, in two different parts of the negative strand. When using a set of randomly permuated starting nucleotides, e.g. TTGTTGTAT, the synthetic sequence generated by Gen-DNA-TCN includes two transcription factor binding sites, i.e. *DAL82* and *RAP1*, in two separated parts of its positive strand. In terms of the transcription factor binding sites in negative strands, the synthetic sequence generated using another permutated starting nucleotides, i.e. CTAATTAAA, includes two transcription factor binding sites, i.e. *LEU3* and *ZAP1*, in two different parts.

**Fig. 10:**
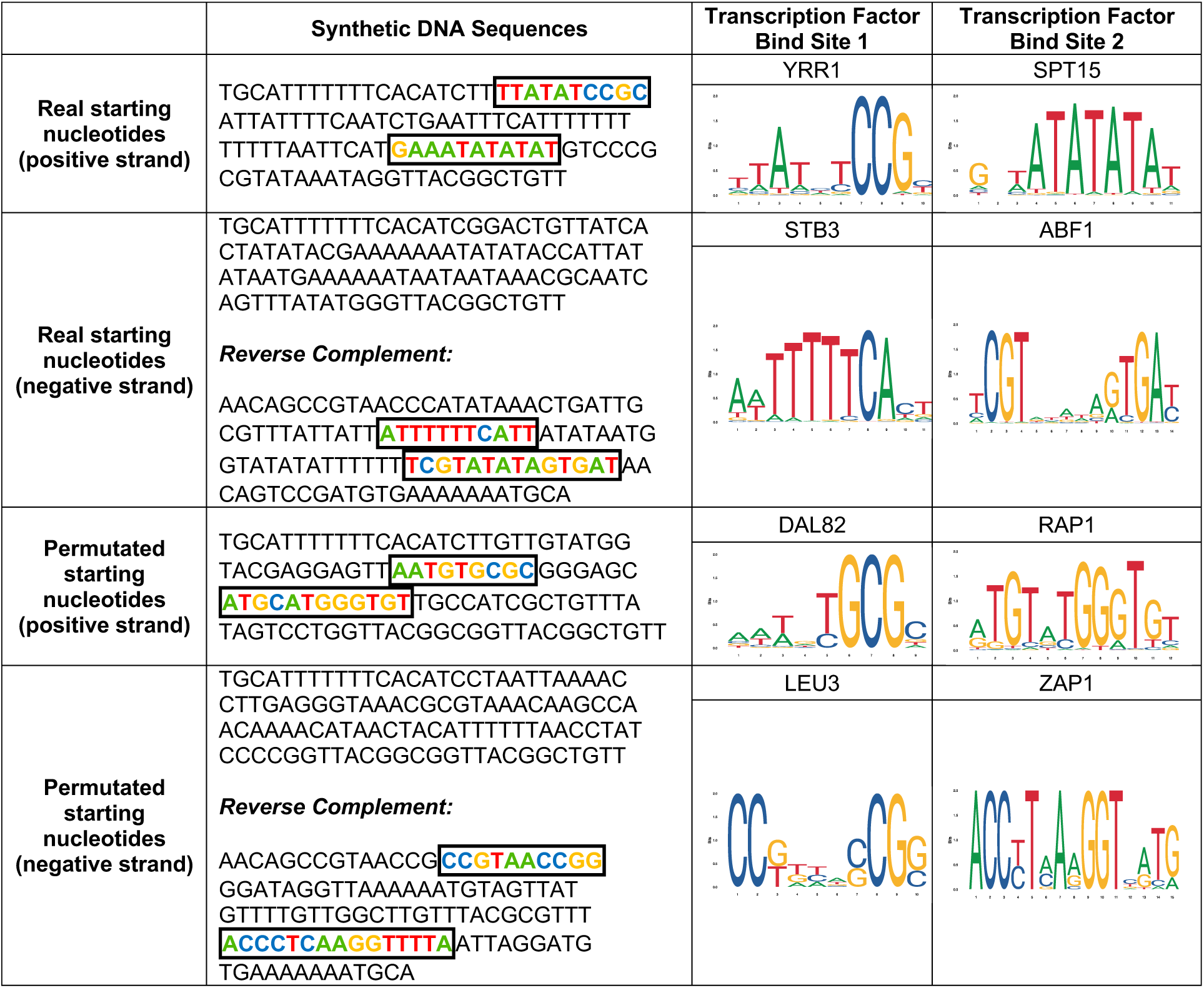
The examples of Gen-DNA-TCN generated synthetic promoter sequences and the detected transcription factor binding site motifs.

### 4.2 The pre-trained Pre-DNA-TCN model facilitates the training of the Gen-DNA-TCN model

As Gen-DNA-TCN exploits the parameters of the pre-trained Pre-DNA-TCN model as a type of prior knowledge for further training, we investigate the role of Pre-DNA-TCN by analysing the training processes of three different models. Figures 11.A - 11.C show the training curves of Model 1, Model 2 and Model 3. In general, the pre-trained Pre-DNA-TCN model using 6,739,258 sequences play a crucial role on the high computational efficiency of the Gen-DNA-TCN training. It is obvious that Model 1 obtained its optimal parameters using the fewest epochs (i.e. 32 epochs), whereas the other two models obtained their corresponding optimal parameters using much more epochs, i.e. 945 epochs for Model 2 and 985 epochs for Model 3. With considering the good performance of Model 1 introduced in previous sections, it is further confirmed that the parameters of the Pre-DNA-TCN model have already encoded sufficient functional sequence information that also facilitates the training of Gen-DNA-TCN, leading to high-quality synthetic sequences and good computational efficiency.

**Fig. 11:**
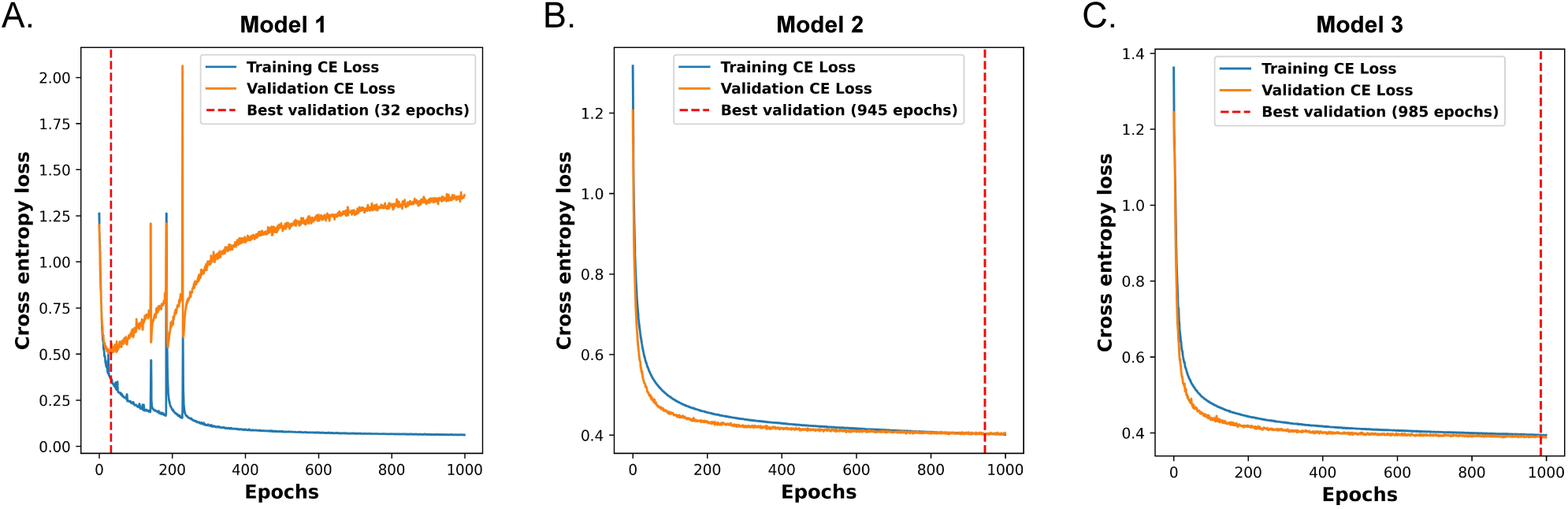
The training curves of three different variants of Gen-DNA-TCN, i.e. Model 1 (A), Model 2 (B) and Model 3 (C).

## 5 Conclusion

In this work, we propose a novel autoregressive generative language model, namely Gen-DNA-TCN, which successfully generates high-quality synthetic functional yeast promoter sequences with exploiting a pre-trained predictive language model encoding functional promoter sequence information. Future research directions would focus on extending the application of the proposed Gen-DNA-TCN model to other synthetic biological sequence generation tasks, e.g. functional protein sequence design.

## References

[1] Rafi, A.M., Nogina, D., Penzar, D., Lee, D., Lee, D., Kim, N., Kim, S., Kim, D., Shin, Y., Kwak, I.-Y., Meshcheryakov, G., Lando, A., Zinkevich, A., Kim, B.-C., Lee, J., Kang, T., Vaishnav, E.D., Yadollahpour, P., Consortium, R.P.D.C., Kim, S., Albrecht, J., Regev, A., Gong, W., Kulakovskiy, I.V., Meyer, P., Boer, C.G.: A community effort to optimize sequence-based deep learning models of gene regulation. Nature Biotechnology 43, 1373–1383 (2025)

[2] Redden, H., Alper, H.S.: The development and characterization of synthetic minimal yeast promoters. Nature Communications 6, 7810 (2015)

[3] Rajkumar, S. Arun, Liu, G., Bergenholm, D., Arsovska, D., Kristensen, M., Nielsen, J., Jensen, M.K., Keasling, J.D.: Engineering of synthetic, stress-responsive yeast promoters. Nucleic Acids Research 44(17), 136 (2016)

[4] Portela, R.M.C., Vogl, T., Kniely, C., Fischer, J.E., Oliveira, R., Glieder, A.: Synthetic core promoters as universal parts for fine-tuning expression in different yeast species. ACS Synthetic Biology 6(3), 471–484 (2017)

[5] Kotopka, B.J., Smolke, C.D.: Model-driven generation of artificial yeast promoters. Nature Communications 11, 2113 (2020)

[6] Zrimec, J., Fu, X., Muhammad, A.S., Skrekas, C., Jauniskis, V., Speicher, N.K., Börlin, C.S., Verendel, V., Chehreghani, M.H., Dubhashi, D., Siewers, V., David, F., Nielsen, J., Zelezniak, A.: Controlling gene expression with deep generative design of regulatory dna. Nature Communications 13, 5099 (2022)

[7] Zhang, P., Wang, H., Xu, H., Wei, L., Liu, L., Hu, Z., Wang, X.: Deep flanking sequence engineering for efficient promoter design using deepseed. Nature Communications 14(6309) (2023)

[8] Gu, Y., Su, J., Xia, J., Wu, P., Wu, H., Su, Y., Wei, P.-J., Zheng, C.-H.: De novo promoter design method based on deep generative and dynamic evolution algorithm. Nucleic Acids Research 53, 833 (2025)

[9] DaSilva, L.F., Senan, S., Kribelbauer-Swietek, J.F.e.a.: Designing synthetic regulatory elements using the generative ai framework dna-diffusion. Nature Genetics (2025)

[10] Oord, A., Dieleman, S., Zen, H., Simonyan, K., Vinyals, O., Alex, G., Kalchbrenner, N., Senior, A., Kavukcuoglu, K.: Wavenet: A generative model for raw audio. arXiv preprint arXiv:1609.03499 (2016)

[11] Bai, S., Kolter, J.Z., Koltun, V.: An empirical evaluation of generic convolutional and recurrent networks for sequence modeling. arXiv preprint arXiv:1803.01271 (2018)

[12] Shin, J.-E., Riesselman, A.J., Kollasch, A.W., McMahon, C., Simon, E., Sander, C., Manglik, A., Kruse, A.C., Marks, D.S.: Protein design and variant prediction using autoregressive generative models. Nature Communications 12, 2403 (2021)

[13] Alsaggaf, I., Greaves, P., Barton, C., Wan, C.: Dream challenge submission report of team. https://github.com/de-Boer-Lab/random-promoter-dream-challenge-2022/tree/main/dream_submissions/Wan%26Barton_BBK (2022)

[14] Khan, A., Fornes, O., Stigliani, A., Gheorghe, M., Castro-Mondragon, J.A., Lee, R., Bessy, A., Chèneby, J., Kulkarni, S.R., Tan, G., Baranasic, D., Arenillas, D.J., Sandelin, A., Vandepoele, K., Lenhard, B., Ballester, B., Wasserman, W.W., Parcy, F., Mathelier, A.: Jasper 2018: update of the open-access database of transcription factor binding profiles and its web framework. Nucleic Acids Research 46(D1), 260–266 (2018)

[15] Grant, C.E., Bailey, T.L., Noble, W.S.: Fimo: scanning for occurrences of a given motif. Bioinformatics 27(7), 1017–1018 (2011)

[16] Bachmann, M.: Levenshtein Distance. https://github.com/rapidfuzz/Levenshtein (2024)

[17] Maaten, L., Hinton, G.: Visualizing high-dimensional data using t-sne. Journal of Machine Learning Research 9, 2579–2605 (2008)

